# Obesity associated with attenuated tissue immune cell responses in COVID-19

**DOI:** 10.1101/2022.01.14.475727

**Authors:** Shuang A. Guo, Georgina S. Bowyer, John R. Ferdinand, Mailis Maes, Zewen K. Tuong, Eleanor Gilman, Mingfeng Liao, Rik G. H. Lindeboom, Masahiro Yoshida, Kaylee Worlock, Huda Gopee, Emily Stephenson, Paul A. Lyons, Kenneth G.C. Smith, Muzlifah Haniffa, Kerstin B. Meyer, Marko Z. Nikolić, Zheng Zhang, Richard G. Wunderink, Alexander V. Misharin, Gordon Dougan, Vilas Navapurkar, Sarah A. Teichmann, Andrew Conway-Morris, Menna R. Clatworthy

## Abstract

Obesity is common and associated with more severe COVID-19, proposed to be in part related to an adipokine-driven pro-inflammatory state. Here we analysed single cell transcriptomes from bronchiolar lavage in three adult cohorts, comparing obese (Ob, body mass index (BMI) >30m^2^) and non-obese (N-Ob, BMI <30m^2^). Surprisingly, we found that Ob subjects had attenuated lung immune/inflammatory responses in SARS-CoV-2 infection, with decreased expression of interferon (IFN)α, IFNγ and tumour necrosis factor (TNF) alpha response gene signatures in almost all lung epithelial and immune cell subsets, and lower expression of *IFNG* and *TNF* in specific lung immune cells. Analysis of peripheral blood immune cells in an independent adult cohort showed a similar, but less marked, reduction in type I IFN and IFNγ response genes, as well as decreased serum IFNα, in Ob patients with SARS-CoV-2. Nasal immune cells from Ob children with COVID-19 also showed reduced enrichment of IFNα and IFNγ response genes. Altogether, these findings show blunted tissue immune responses in Ob COVID-19 patients, with clinical implications.

## Main

SARS-CoV-2 is a novel coronavirus responsible for the current global pandemic, with more than 275 million cases and 5.3 million deaths confirmed worldwide (World Health Organisation, Dec 24^th^ 2021). The clinical course of SARS-CoV-2 is variable, ranging from asymptomatic disease to acute respiratory distress syndrome requiring ventilatory support^1^. Those at risk of a more severe clinical course following infection include the elderly, immunosuppressed, and those with co-morbidities including obesity^1–4^. Despite the expedited delivery of clinically validated SARS-CoV-2 vaccines, some ‘vulnerable’ groups mount poor vaccine responses, and emerging viral variants are poorly neutralised by vaccine-induced antibodies^5^. Thus, there is an on-going need to better understand viral immune responses in those at highest risk of severe disease, such as obese individuals.

Obesity, defined as a body mass index (BMI) of >30m^2^, is common and affects more than 40% of US adults^6^. Whilst an increased BMI generates mechanical factors that may compromise ventilation^7^, obesity is also associated with an inflammatory state characterised by elevated circulating cytokines such as tumour necrosis factor (TNF) and interleukin (IL) 6^8^, chemokines, including the neutrophil recruiting chemokine CXCL8^9^, and the monocyte chemoattractant CCL2 (MCP1) that mediates the accumulation of inflammatory adipose tissue macrophages^10^. Lymphocyte abnormalities have also been noted in obesity, and an increased frequency of circulating interferon (IFN)-γ secreting CD4 T cells noted, the latter likely related to the known effects of leptin in promoting Th1 polarisation^11^. Indeed, the appetite-regulating hormone leptin, produced by adipocytes and increased in obesity, has several direct immune stimulatory effects, promoting NK cell cytotoxicity, antigen presentation by dendritic cells (DC) and monocyte and B cell secretion of TNF and IL6 by engagement of the leptin receptor (LEPR) which is expressed on many immune cells, and principally acts by triggering JAK2/STAT3 signalling^12^. Leptin levels increase in lean animals challenged with pro-inflammatory cytokines, infectious agents or pathogen-derived molecules^13^, and in COVID-19, elevated serum leptin levels have been described in ventilated SARS-CoV-2 patients compared with controls^14^, with a positive correlation with BMI^15^.

There has been an intense focus on delineating the nature of the immune response in SARS-CoV-2 infection; Type I IFN responses play a critical role in protective immunity, with genetic-deficiency or neutralising autoantibodies affecting this axis mediating increased susceptibility to severe infection^16^. Transcriptomic studies have identified both absent/low and increased IFN response genes in patients with severe or lethal disease, and longitudinal studies revealed a diminished and/or delayed induction of type I IFNs in COVID-19 patients compared with patients with influenza, with an exuberant early TNF/IL6 response^16^. To date, the effect of obesity on immune responses to SARS-CoV-2, particularly tissue responses, has not been considered, although it has been proposed that the elevated leptin associated with obesity might promote an excessive inflammatory response, contributing to the worse outcomes observed in obese patients with COVID-19, with calls for anti-inflammatory therapeutic strategies to be employed in this patient group^17^.

Here we profiled paired blood and bronchoalveolar lavage (BAL) samples from 4 patients with severe COVID-19 requiring mechanical ventilation and intensive care treatment, and 4 control ventilated non-COVID-19 patients using flow cytometry and scRNAseq. To address the issue of whether patients with a high BMI have abnormal tissue immune responses to SARS-CoV-2, we integrated our scRNAseq data (UCAM) with two previous COVID-19 BAL scRNAseq datasets from Shenzhen 3^rd^ Hospital, China (SZH) and Northwestern University, Chicago, USA (NU)^18,19^ and obtained the BMI metadata associated with these samples, enabling a comparison of BAL immune cells in 13 obese (Ob) patients (BMI>30) and 20 non-obese (N-Ob) (BMI<30) COVID-19 patients and ventilated non-COVID controls (**Fig. 1a, S1a**). Overall, Ob subjects were more prevalent in the NU cohort compared to the other two cohorts and underwent BAL sampling at earlier time-points following admission to the intensive care unit (**Fig. S1a**).

**Figure 1.**
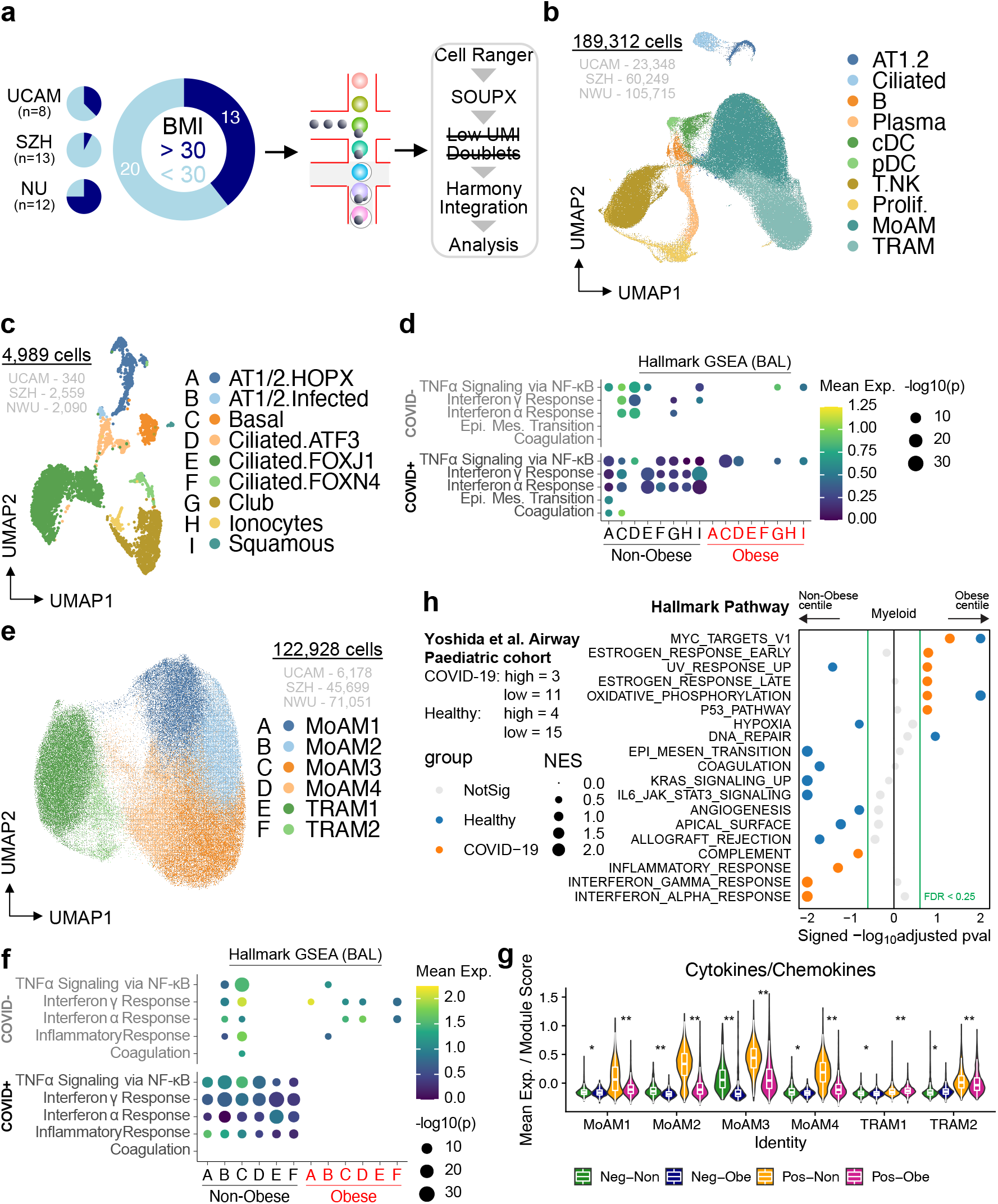
Single-cell analysis of bronchoalveolar lavage (BAL) fluid samples from patients with or without COVID-19 reveals differences in geneset enrichment in structural and myeloid cells in non-obese compared with obese subjects. **a.** Overview of the workflow. In this study, we included BAL samples from 33 patients from three cohorts with BMI information, namely UCAM (n=8; ob=3; this study), SZH (n=13; ob=1; Liao et al., 2020), and NU (n=12; ob=9; Grant et al., 2021). b. UMAP embedding of 189,312 cells post-integration of the 3 datasets. Cells are coloured according to harmonised broad cell type annotations. c. UMAP embedding of 4,989 epithelial and structural cells post-integration coloured according to harmonised fine cell type annotations. d. Dot plot of gene set enrichment analysis of top 5 most enriched immune pathways within Hallmark gene sets for epithelial/structural cells. Mean expression of genes contained in each gene set within each cell type (labelled A-I; see c.), separated into non-obese vs. obese groups, are indicated by colour gradients. P values are indicated by dot sizes. e. UMAP embedding of 122,928 myeloid cells coloured according to fine cell type annotation as per Grant et al. (2021). MoAM indicates monocyte-derived macrophage, TRAM indicates tissue-resident macrophage f. Dot plot of gene set enrichment analysis of top 5 most enriched immune pathways within Hallmark gene sets for myeloid cells. Mean expression of genes contained in each gene set within each cell type (labelled A-F; see e.), separated into non-obese vs. obese groups, are indicated by colour gradients. P values are indicated by dot sizes. g. Violin plot depicting mean expression levels of transcripts for cytokines and chemokines in each macrophage subpopulation. Differences between Non-obese vs. Obese with or without COVID-19 infection remain significant by Wilcoxon rank sum test. h. Dot pot of gene set enrichment analysis of Hallmark gene sets in myeloid cells from paediatric airway samples (Yoshida et al. (2021)) between non-obese vs. obese children. Normalized enrichment score (NES) of each pathway is indicated by dot size. Colour of circles indicate which comparison was significantly enriched (healthy or COVID-19); grey circles are not significant.

Following QC, dataset integration and batch correction, data was available on 189,312 cells. Clusters were broadly annotated using canonical marker expression and comparison to previously published BAL single cell datasets, to identify alveolar type 1 and 2 pneumocytes (AT1/2), ciliated cells, B and plasma cells, classical and plasmacytoid dendritic cells (c/pDCs), a broad T cell/innate lymphocyte cluster and alveolar macrophages, including monocyte-derived and tissue-resident clusters, with reasonable representation of cells from Ob and N-Ob patients in all clusters (**Fig. 1b, S1b-c**).

Analysis of the non-immune cells in isolation enabled the identification of several ciliated epithelial cell subsets, as well as basal cells, club cells, squamous cells and alveolar type 1 and AT1/2 (**Fig. 1c, S2a**). Organ structural cells may contribute to tissue immune responses, and indeed, several immune-related transcripts were highly expressed in some of these subsets; SARS-CoV-2-infected AT1/2 cells expressed neutrophil recruiting chemokine transcripts *(CXCL1, CXCL2* and *CXCL8),* but this was attenuated in Ob compared with N-Ob subjects (**Fig. S2b**). Consistent with this, flow cytometric analysis of the UCAM cohort BAL confirmed a reduction in neutrophils in Ob BAL compared with N-Ob BAL (**Fig. S2c**). *IL1RN* (encoding IL1RA, a protein that binds to IL1R inhibiting the pro-inflammatory effects of IL1β, including arterial inflammation^20^) was highly expressed in squamous epithelial cells in N-Ob, whilst barely detectably in Ob subjects (**Fig. S2b**). Conversely, GDF15 the pro-cachectic cytokine^21^ was more highly expressed in Ob subjects across a range of airway cells (**Fig. S2b**). Surprisingly, given the association of obesity with inflammation, gene-set enrichment analysis (GSEA) demonstrated a marked negative enrichment of *interferon-alpha* and/or *interferon-gamma response genes* in all non-immune cell clusters in Ob compared with N-Ob COVID-19+ BAL, with more variable differences in these genesets in COVID-19 negative patients (**Fig. 1d**).

Considering alveolar macrophages in isolation, four subsets of monocyte-derived macrophages (MoAM), and two subsets of tissue resident (TRAM1/2) macrophages were evident, as noted previously^18,19^ (**Fig. 1e, S2d**). In Ob subjects with SARS-CoV-2 infection, all alveolar macrophage subsets showed reduced enrichment of *interferon-alpha* and/or *interferon-gamma response genes, Tnfa via NFkB, Inflammatory response,* and *complement* pathway genes, with reduced *JAK-STAT3 signalling pathway* genes also evident in MoAM1,2 and 4 compared with N-Ob COVID-19 (**Fig. 1f, S2e**), with a similar pattern observed in cDCs and pDCs in BAL (**Fig.S2f**). Indeed, a curated chemokine/cytokine gene module score was significantly higher in N-Ob alveolar macrophages, particularly in monocyte-derived AM subsets (**Fig. 1g**). Given the differences in timing of BAL sampling between groups, we plotted individual patient gene pathway enrichment scores against time (**Fig. S2g**). This confirmed that *interferon-alpha* and/or *interferon-gamma response* genes were reduced in Ob compared with N-Ob subjects, across timepoints, and in all AM subsets except TRAM2, which showed only an early attenuation of responses in Ob (**Fig. S2g**). Genes attenuated in Ob alveolar macrophages and cDCs included *CXCL10,* a classical IFNγ response gene previously proposed to form a pro-inflammatory circuit with T cells^19^, and several monocyte and lymphocyte-recruiting chemokines (**Fig. S2h-i**). A notable exception to the muted cytokine and chemokine expression in Ob macrophages, cDCs and pDCs was *TGFB1,* a tissue repair factor, which in excess can contribute to fibrosis, which was broadly more highly expressed in Ob cells in COVID-19 (**Fig. S2h-i**). To validate these findings and confirm their relevance across lifespan, we examined the single cell transcriptomes of tissue immune cells isolated from nasal brushings in children with SARS-CoV-2^22^. We similarly found reduced enrichment of *interferon-alpha* and/or *interferon-gamma response* genes in myeloid cells in Ob children compared with N-Ob (**Fig. 1h**).

We next assessed T cells and innate lymphocytes in adult BAL, annotating naïve and effector memory CD4 and CD8 subsets, Tregs, MAIT cells, NKT and two subsets of NK cells, CD56^high^ and CD56^low^ (**Fig. 2a, S3a**). GSEA again demonstrated reduced enrichment of *interferon-alpha* and/or *interferon-gamma response* genes*, and Tnfa signalling via NFkB,* pathway genes across every subset present in Ob BAL (**Fig. 2b, S3b-c**). In Ob children with SARS-CoV-2, nasal T cells and innate lymphocytes cells also showed reduced enrichment of *interferon-alpha* and/or *interferon-gamma response* genes compared with N-Ob (**Fig. 2c, S3d**).

**Figure 2.**
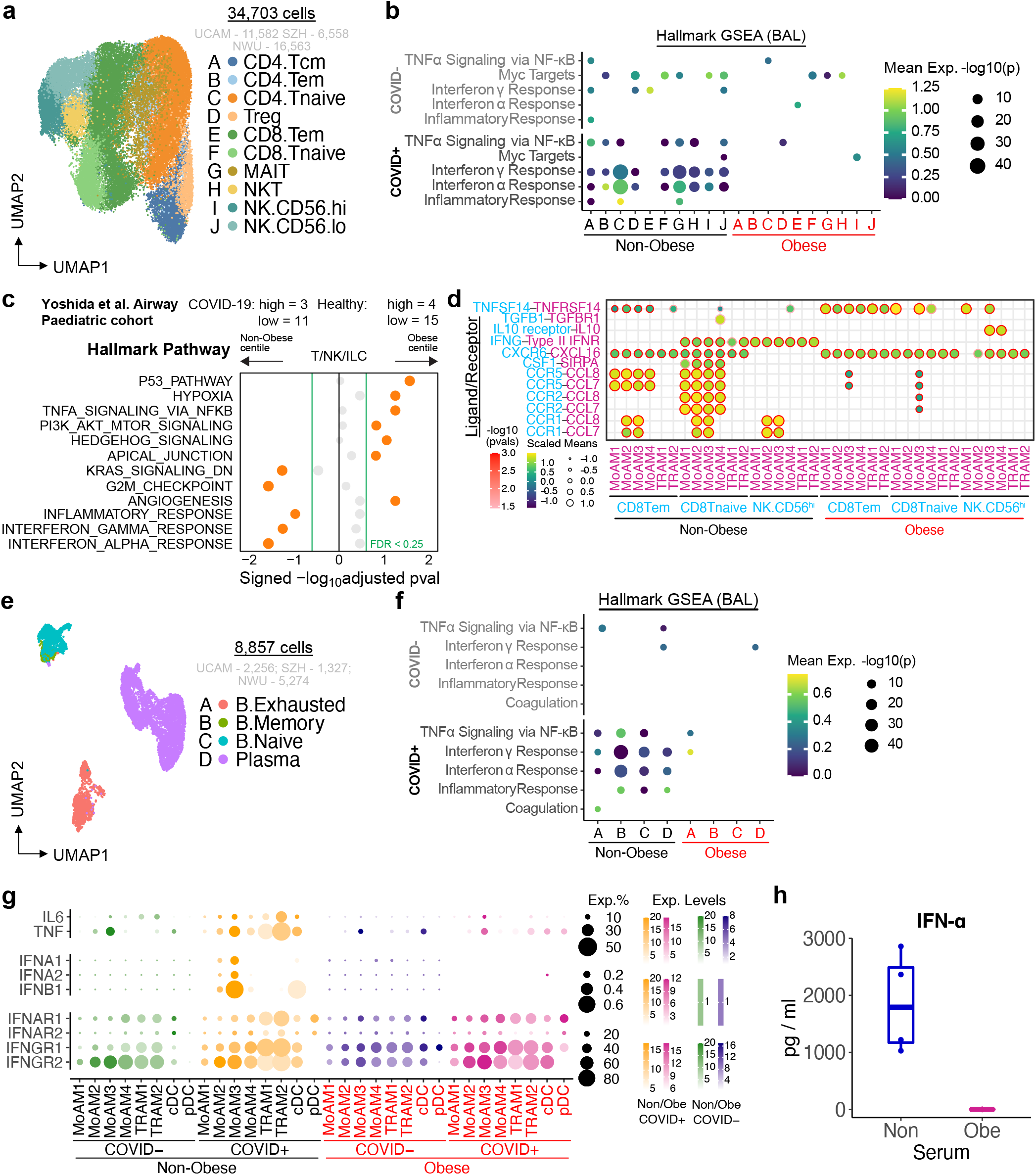
Single-cell analysis of lymphocytes shows reduced enrichment of type I and gamma interferon response genes in obese COVID-19 patients. a. UMAP embedding of 34,703 T/NK cells post-integration. CD4 T cells (Tcm: central memory T cells; Tem: effector memory T cells; Tnaive: naive T cells; and Treg), CD8 T cells (Tem, Tnaive, and MAIT), NKT cells, and NK cells (CD56-high or CD56-low). b. Dot plot of gene set enrichment analysis of top 5 most enriched immune pathways within Hallmark gene sets for T/NK cells. Mean expression of genes contained in each gene set within each cell type (labelled A-J; see a.), separated into non-obese vs obese groups, are indicated by colour gradients. P values are indicated by dot sizes. c. Dot pot of gene set enrichment analysis of Hallmark gene sets in T/NK/ILC cells from paediatric airway samples (Yoshida et al. (2021)) between non-obese vs. obese children. Normalized enrichment score (NES) of each pathway is indicated by dot size. Colour of circles indicate which comparison was significantly enriched (healthy or COVID-19); grey circles are not significant. d. Ligand-receptor analysis with CellPhoneDB infers distinct interactions between CD8.Tem / CD8.Tnaive / NK.CD56hi and alveolar macrophages. Size and colour gradient of circles indicate the scaled interaction score; interaction scores are scaled row-wise. Red outline indicates a P value < 0.05. e. UMAP embedding of 8,857 B/plasma cells post-integration coloured according to harmonised fine cell type annotations. f. Dot plot of gene set enrichment analysis of top 5 most enriched immune pathways within Hallmark gene sets for B/plasma cells. Mean expression of genes contained in each gene set within each cell type (labelled A-J; see e.), separated into non-obese vs. obese groups, are indicated by colour gradients. P values are indicated by dot sizes. g. Mean expression dot plots of transcripts for IFN-α/β/γ, IFN receptors, IL6, and TNF-α in myeloid cells in the BAL samples. Expression levels in each case are indicated by distinct colour gradients (Green: Non-obese without COVID-19; Yellow: Non-obese with COVID-19; Purple: Obese without COVID-19; Magenta: Obese with COVID-19). Expression percentages are indicated by dot sizes. h. Serum IFN-α measurements from n=4 obese and n=4 Non-obese patients from the adult PBMC patient cohort (Stephenson et al.). All 4 Obese samples were below detection limits.

In adult BAL, predictive cell-cell interaction analysis based on receptor-ligand expression suggested reduced alveolar macrophage production of chemokines *(CCL7, CCL8)* predicted to attract CD8 T and NK cell subsets in Ob subjects (**Fig. 2d**), both important for anti-viral immunity. Few predicted interactions were increased in Ob patients, notably IL10-IL10R (MoMAC3/4 and NKCD56^hi^) with the potential to suppress NK cell cytotoxicity^23^, and TNFSF14-TNFRSF14 (MoMAC1/3 and naïve CD8 T cells) (**Fig. 2d**), encoding LIGHT-HVEM, an axis important for stimulating lymphocytes, but high serum LIGHT levels have been associated with fatal COVID-19^24^.

B cell and plasma cells clusters in adult BAL included naïve B cells, memory B cells, exhausted B cells and plasma cells. (**Fig. 2e, S4a-b**). In COVID-19, there was again reduced enrichment of *interferon-alpha* and/or *interferon-gamma response* and *Tnfa signalling via NFkB* pathway genes in all BAL B cell subsets in Ob subjects, with more variable effects in the paediatric samples (**Fig. 2f, S4c-d**).

To determine if blunted immune responses were evident in Ob subjects beyond tissue immune cells, we obtained BMI data (where available) on an additional cohort of adult COVID-19 patients recently included in a multiomic analysis of PBMCs^25^. In this cohort, patient’s blood was sampled between day 0 and 20 post-symptom onset, with a similar temporal distribution of sampling in Ob and Non-Ob (**Fig. S5a**). As observed in adult BAL, there was reduced enrichment of *interferon-alpha* and/or *interferon-gamma response genes* in peripheral blood T cells, innate lymphocytes and B cells in Ob COVID-19 patients compared with N-Ob (**Fig. S5b-c**). Interestingly, *Tnfa signalling via NFkB* pathway genes showed the opposite enrichment pattern to that observed in BAL, with an increase in Ob COVID-19 patients (**Fig. S5b-c**).

We reasoned that the reduced tissue cell responses to IFNα, IFNγ and TNFa might be due to a decreased production of these cytokines, or a reduced ability to respond to them due to decreased receptor expression. To distinguish between these possibilities, we first assessed cytokine transcripts in BAL cells; *IFNA1/2* and *IFNB* transcripts were undetectable, except in <0.5% of MoAM3 in N-Ob COVID-19+ subjects (**Fig.2g, S6**). There was little expression of type I IFN transcripts in peripheral blood immune cells (**Fig. S7**), but serum cytokine measurement showed undetectable IFNα levels in Ob but elevated levels in N-Ob subjects (**Fig. 2h**), with no significant differences in TNF (which was undetectable) and IFNγ (data not shown). *IFNAR1/2* expression was not decreased in Ob versus N-Ob COVID-19 BAL cells, with an increase in *IFNAR1* in some alveolar macrophage subsets in Ob COVID-19 (**Fig. 2g, S6**). Altogether, this is consistent with a failure of type I IFN production rather than a reduced ability to respond to type I IFNs in Ob subjects in COVID-19. In SARS-CoV-2 infected patients, *IFNG* transcripts were detectable in proliferating lymphocytes, CD4 central memory T cells, naïve CD8 T cells and NKT cells in BAL, again mainly at a higher level in N-Ob compared with Ob patients (**Fig. S6**). In BAL, *TNF* transcripts were highest in N-Ob COVID-19+ MoAM3, TRAM2, and cDC (**Fig. 2g, S6**), but in blood, the opposite was observed, with higher *TNF* and lower *TNFRSF1A/B* expression in some subsets of Ob circulating monocytes (**Fig. S8**).

Obese patients are known to have basal immune activation and inflammation, in part due to the immunostimulatory effects of elevated leptin. It has been proposed that this contributes to the increased susceptibility to severe SARS-CoV-2 infection and the worse outcomes observed in obese subjects, generating a heightened pro-inflammatory cytokine response. In fact, our analysis of Ob adult BAL and Ob paediatric nasal immune cells in COVID-19 suggests that these patients exhibit a broadly immunosuppressed state in tissues compared with non-obese subjects, with reduced type I IFN and IFN gamma signatures across almost all immune cell subsets, as well as decreased expression of monocyte and neutrophil recruiting chemokines. Of note, our findings bare similarity to studies of obese mice challenged with influenza, which not only had increased mortality, but decreased *Ifna, Ifnb* and *Ifng* transcripts in lung tissue, as well as lower levels of some chemokines *(Ccl2* and *Ccl5),* compared with lean controls, despite a higher viral load ^13,26^. In addition, these obese animals also had impaired antigen presentation by DC, decreased IFN-γ production by memory T cells, and reduced NK cell cytotoxicity in this model^27^. Interestingly, serum leptin concentrations increased in lean mice during influenza infection, in contrast to obese mice, where leptin decreased, such that during infection levels were similar to lean mice^13^ but leptin resistance in the latter contributing to attenuated immune cell responses^28^. Consistent with this, in COVID-19 BAL, we observed reduced *JAK-STAT3 signalling* pathway genes in a number of lung immune cell subsets in Ob subjects, particularly in monocyte-derived alveolar macrophages.

Notably, although IFN response genes were reduced in Ob subjects in peripheral blood, this phenomenon was much less marked outside of tissues and there was a disconnect between BAL and blood in terms of TNF response genes, suggesting attenuated tissue responses, but a more exuberant, potentially pathogenic systemic pro-inflammatory landscape, and emphasising the importance of studies assessing tissue immunity, despite the practical challenges associated.

Overall, our study has important translational implications; current and proposed treatments for severe COVID-19 include anti-inflammatory agents such as IL6R blocking antibodies and the application of recombinant IFNα and IFNβ to promote early anti-viral responses, with the latter showing limited efficacy in an early trial^29^. However, this study was small and included only patients with mild disease, and there is an increasing recognition for the need to tailor the different treatment strategies available to the correct patient group, at the correct time^16^. Our data show a markedly muted response to type I IFN and IFNγ in tissue immune cells in the respiratory tract in Ob COVID-19 patients across lifespan, as well as reduced transcripts of these cytokines, supporting the application of locally-delivered, inhaled recombinant type I IFNs to respiratory tract tissues in this vulnerable subset.

## Acknowledgements

GB is funded by a Wellcome Strategic Scientific award (WT211276/Z/18/Z). ZKT and MRC are supported by a Medical Research Council Research Project Grant (MR/S035842/1). JRF and MRC are supported by the National Institute of Health Research (NIHR) Blood and Transplant Research Unit in Organ Donation, and NR, MM, GD and MRC by the NIHR Cambridge Biomedical Research Centre. MZN acknowledges funding from the Rutherford Fund Fellowship allocated by the Medical Research Council and the UK Regenerative Medicine Platforms 2 (MR/5005579/1). KBM acknowledges funding from Wellcome (WT211276/Z/18/Z and Sanger core grant WT206194), the Chan Zuckerberg Foundation (grants 2017-174169 and 2019-202654) and the European Union’s Horizon 2020 research and innovation programme under grant agreement No 874656. The views expressed are those of the author(s) and not necessarily those of the NHS, the NIHR or the Department of Health and Social Care. The CL3 for this research was partly funded by the NIHR AMR Research Capital Funding Scheme [NIHR200640]. We are grateful to the Evelyn Trust (20/75), Addenbrooke’s Charitable Trust, Cambridge University Hospitals (12/20A), the NIHR Cambridge Biomedical Research Centre, Rosetrees Trust (M944), Action Medical Research (GN2911) and the UKRI/NIHR through the UK Coronavirus Immunology Consortium (UK-CIC) for their financial support. RGW and AVM are funded by NIH NIAID U19AI35964. ACM is supported by a Clinician Scientist Fellowship from the Medical Research Council (MR/V006118/1). MRC and SAG by an NIHR Research Professorship RP-2017-08-ST2-002).

---

**Figure S1.**
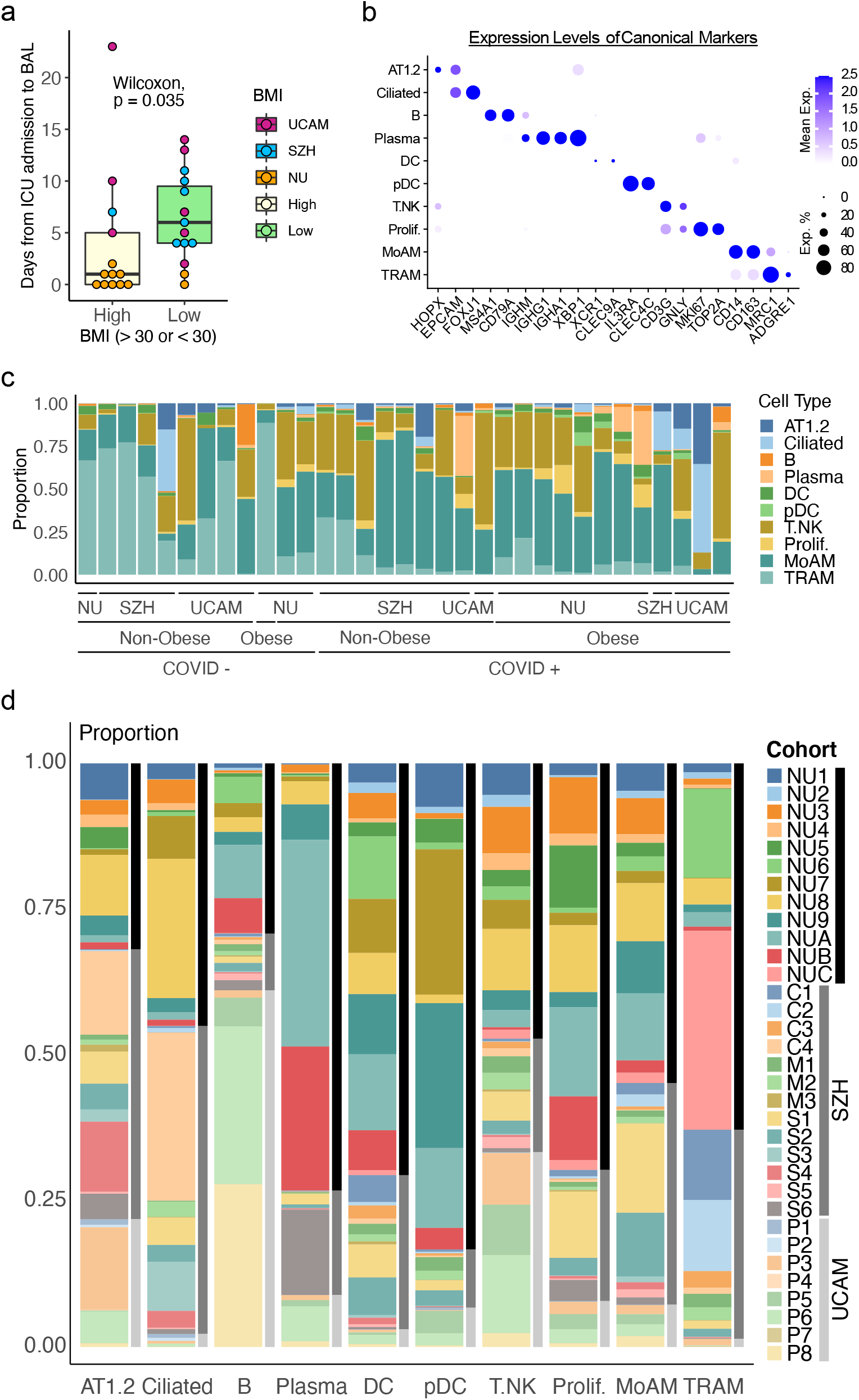
Patient sampling and cellular composition of BAL. a. Distribution of patient’s days from ICU admission to BAL sampling between obese (BMI>30) and non-obese (BMI<30) groups. Wilcoxon rank sum test where p <0.05 was considered statistically significant. b. Mean expression dot plot of transcripts for canonical marker genes in each major population in the BAL samples. Expression levels are indicated by colour gradients. Expression percentages are indicated by dot sizes. c. Proportions of each major population in each cohort grouped by infection (COVID+/-) and obesity (Non-Obese/Obese) states. Each bar indicates the cell type proportions in an individual sample. d. Proportions of each major population in each cohort contributed by each sample (indicated by colour codes) (Black: NU; Gray: SZH; Light Gray: UCAM). Each bar indicates a cell type.

**Figure S2.**
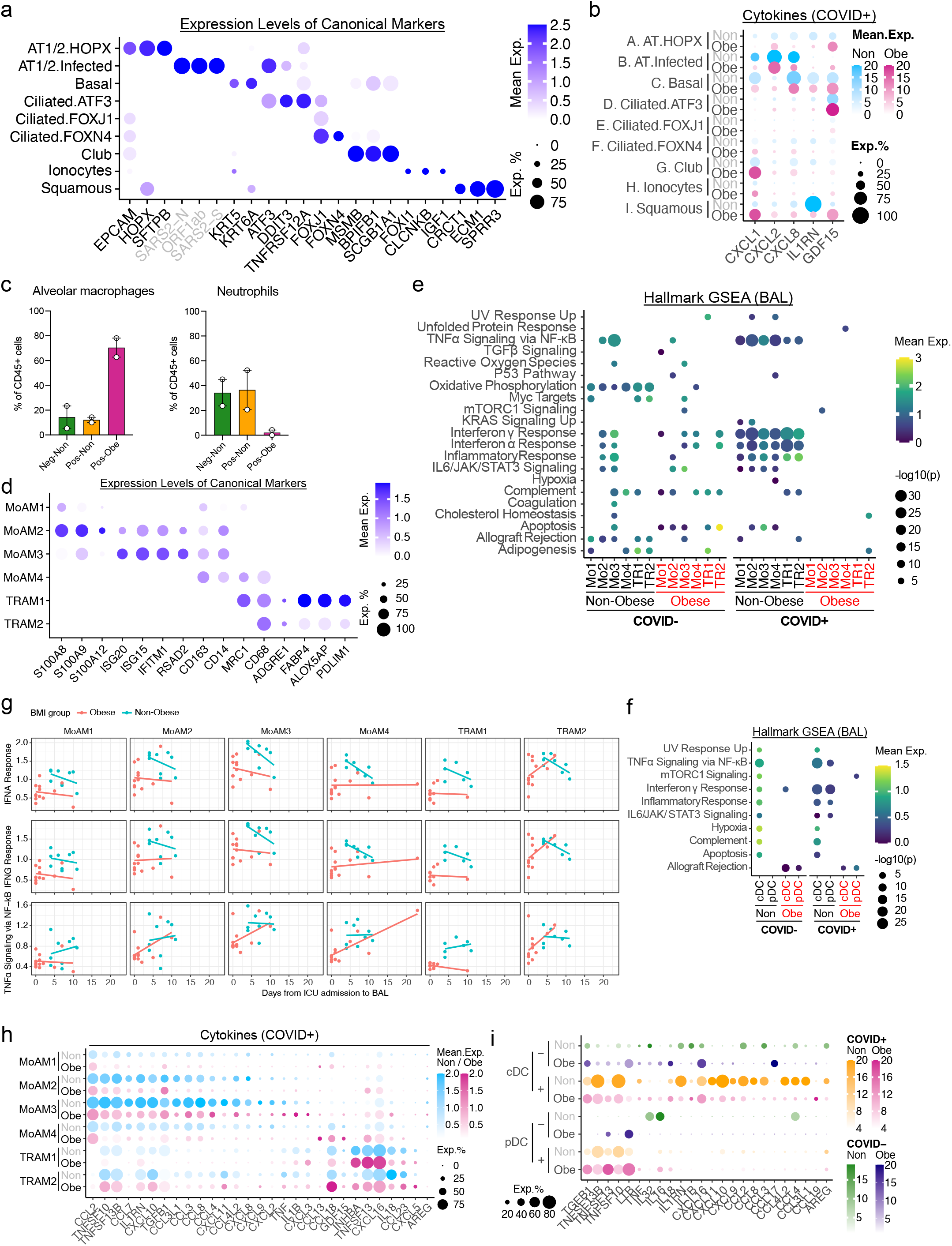
Structural cells and myeloid cells in the BAL samples. a. Mean expression dot plot of transcripts for canonical marker genes in each epithelial subpopulation in the BAL samples. Expression levels are indicated by colour gradients. Expression percentages are indicated by dot sizes. b. Mean expression dot plot of the top differentially expressed cytokines and chemokines in each epithelial subpopulation in the COVID+ BAL samples. Expression levels in each case are indicated by distinct colour gradients (Blue: Non-obese; Magenta: Obese). Expression percentages are indicated by dot sizes. c. Flow cytometry quantification of alveolar macrophages (SSC^hi^CD206^+^, left) and neutrophils (SSC^hi^CD206^-^CD24^+^CD16^+^, right) as a proportion of live CD45^+^ cells in the BAL fluid of obese and non-obese SARS-CoV-2 positive/negative patients (n=6 patients from the Cambridge study only). d. Mean expression dot plot of transcripts for canonical marker genes in each macrophage subpopulation in the BAL samples. Expression levels are indicated by colour gradients. Expression percentages are indicated by dot sizes. MoAM: Monocyte-derived alveolar macrophage; TRAM: Tissue-resident alveolar macrophage. e. Dot plot of gene set enrichment analysis of most enriched immune pathways within Hallmark gene sets for each macrophage subpopulation. Mean expression of genes contained in each gene set within each cell type, separated into non-obese vs. obese and COVID-vs. COVID+ groups, are indicated by colour gradients. P values are indicated by dot sizes. f. Dot plot of gene set enrichment analysis of most enriched immune pathways within Hallmark gene sets for classical dendritic cells (cDC) and plasmacytoid dendritic cells (pDC). Mean expression of genes contained in each gene set within each cell type, separated into non-obese vs. obese and COVID-vs. COVID+ groups, are indicated by colour gradients. P values are indicated by dot sizes. g. Scatter plot of mean expression levels (y-axis) of the leading-edge genes in the signaling pathways IFNα Response, IFNγ Response, and TNFα Signaling via NF-kB versus days from ICU admission to BAL sampling (x-axis) across macrophage subpopulations. MoAM, monocyte-derived alveolar macrophage; TRAM, tissue-resident alveolar macrophage. h. Mean expression dot plot of the top differentially expressed cytokines and chemokines in each macrophage subpopulation in the COVID+ BAL samples. Expression levels in each case are indicated by distinct colour gradients (Blue: Non-obese; Magenta: Obese). Expression percentages are indicated by dot sizes. i. Mean expression dot plots of transcripts for the top differentially expressed cytokines and chemokines in each dendritic cell subpopulation in the BAL samples. Expression levels in each case are indicated by distinct colour gradients (Green: Non-obese without COVID-19; Yellow: Non-obese with COVID-19; Purple: Obese without COVID-19; Magenta: Obese with COVID-19). Expression percentages are indicated by dot sizes.

**Figure S3.**
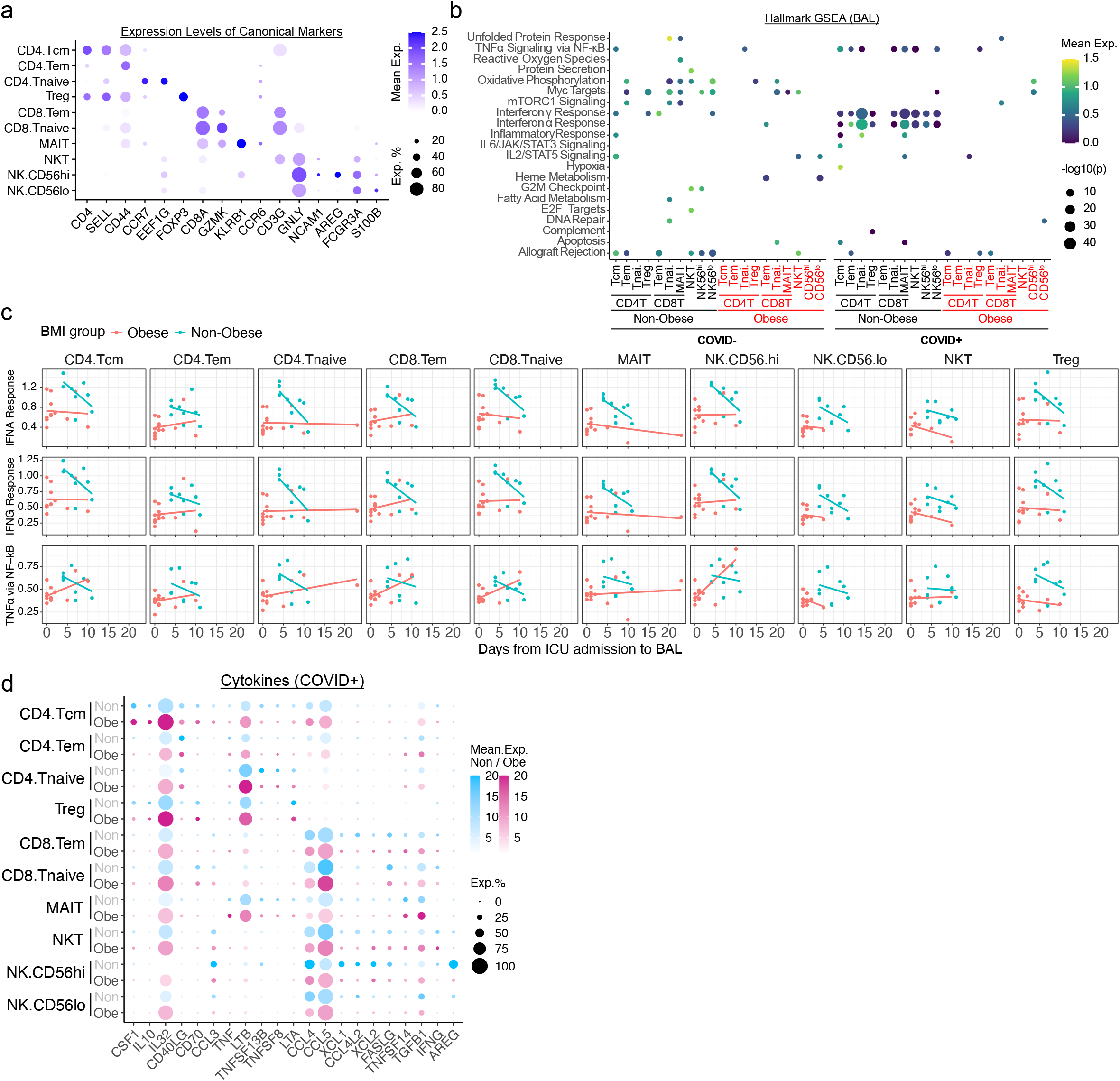
T cells and NK cells in the BAL samples. a. Mean expression dot plot of transcripts for canonical marker genes in each T/NK subpopulation in the BAL samples. Expression levels are indicated by colour gradients. Expression percentages are indicated by dot sizes. Tcm: central memory T cells; Tem: effector memory T cells; Tnaive: naive T cells; Treg: regulatory T cells; MAIT: mucosal associated invariant T cells. b. Dot plot of gene set enrichment analysis of most enriched immune pathways within Hallmark gene sets for each T/NK subpopulation. Mean expression of genes contained in each gene set within each cell type, separated into non-obese vs. obese and COVID-vs. COVID+ groups, are indicated by colour gradients. P values are indicated by dot sizes. c. Scatter plot of mean expression levels (y-axis) of the leading-edge genes in the signaling pathways IFNα Response, IFNγ Response, and TNFα Signaling via NF-kB versus days from ICU admission to BAL sampling (x-axis) across T or NK subpopulations. d. Mean expression dot plot of the top differentially expressed cytokines and chemokines in each T/NK subpopulation in the COVID+ BAL samples. Expression levels in each case are indicated by distinct colour gradients (Blue: Non-obese; Magenta: Obese). Expression percentages are indicated by dot sizes.

**Figure S4.**
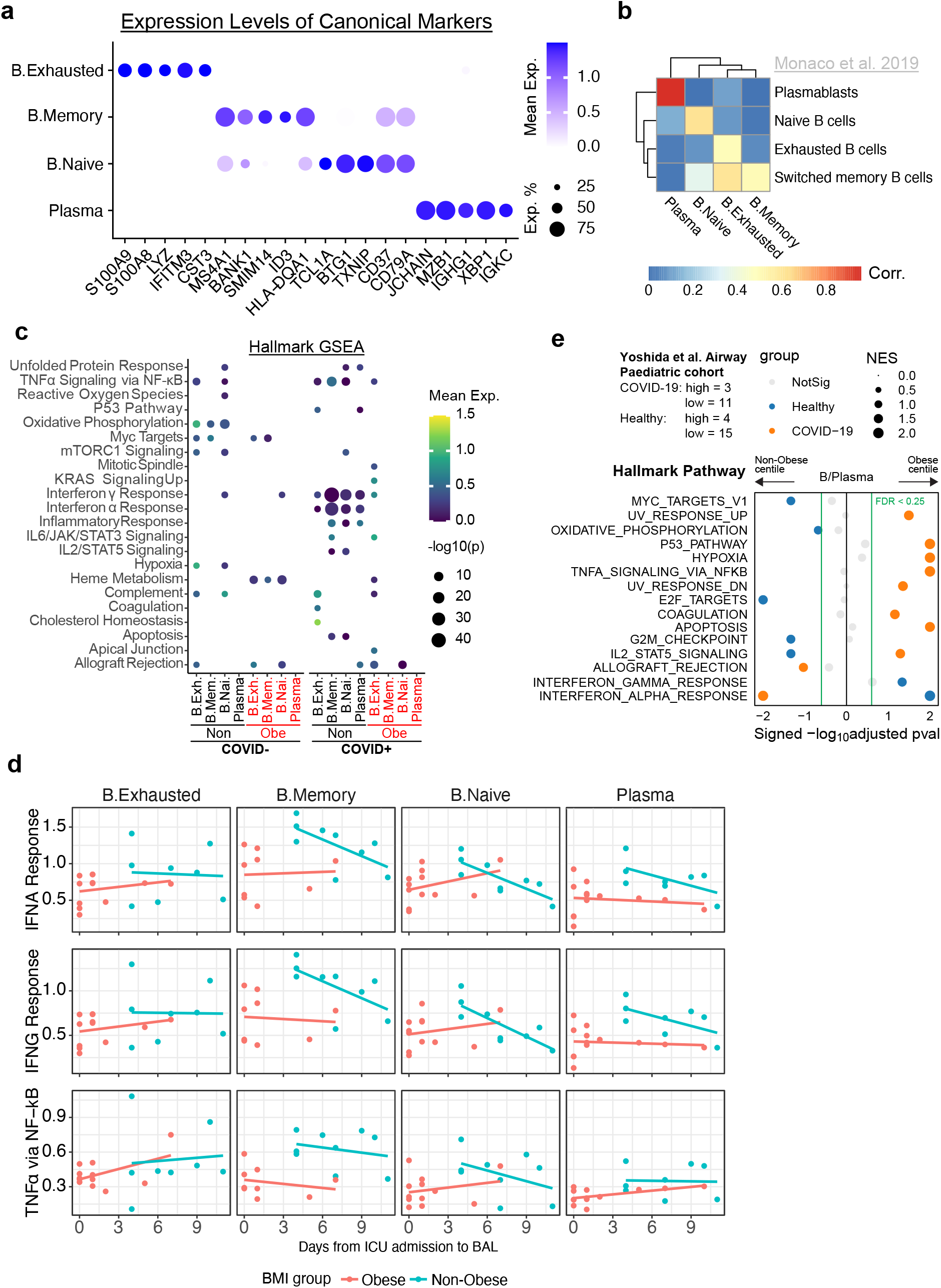
B cells and plasma cells in the BAL samples. a. Mean expression dot plot of transcripts for canonical marker genes in each B/Plasma subpopulation in the BAL samples. Expression levels are indicated by colour gradients. Expression percentages are indicated by dot sizes. b. Heatmap of Pearson’s correlation analysis between our annotated cell types and the public data from Monaco et al. 2019 c. Dot plot of gene set enrichment analysis of most enriched immune pathways within Hallmark gene sets for each B/Plasma subpopulation. Mean expression of genes contained in each gene set within each cell type, separated into non-obese vs. obese and COVID-vs. COVID+ groups, are indicated by colour gradients. P values are indicated by dot sizes. d. Scatter plot of mean expression levels (y-axis) of the leading-edge genes in the signaling pathways IFNα Response, IFNγ Response, and TNFα Signaling via NF-kB versus days from ICU admission to BAL sampling (x-axis) across B subpopulations and plasma cells. e. Dot pot of gene set enrichment analysis of Hallmark gene sets in B/Plasma cells from paediatric airway samples (Yoshida et al. (2021)) between non-obese vs. obese children. Normalized enrichment score (NES) of each pathway is indicated by dot size. Colour of circles indicate which comparison was significantly enriched (healthy or COVID-19); grey circles are not significant.

**Figure S5.**
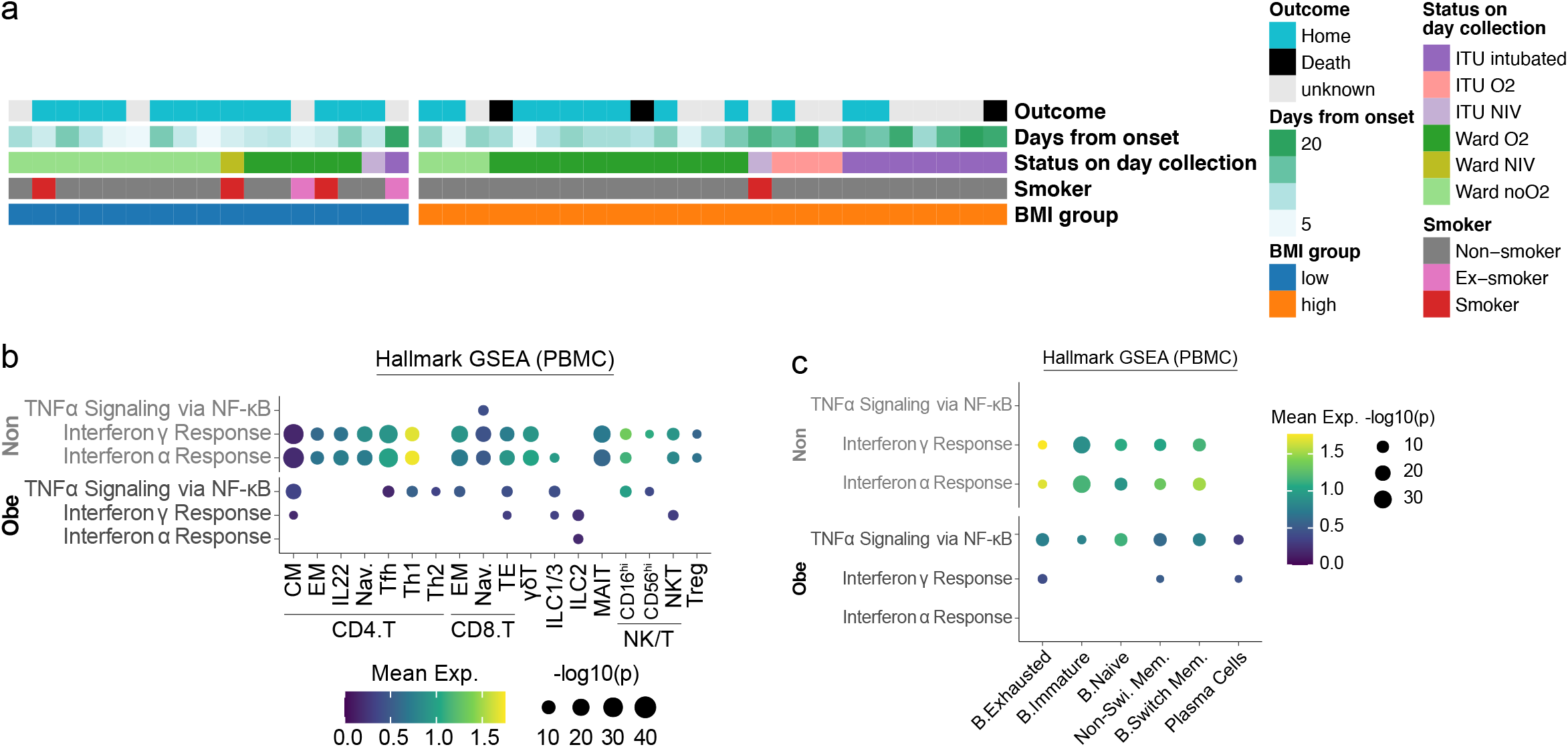
Validation of the signaling pathways in PBMC samples. a. Sample demographics of Stephenson et al. 2021 ^22^ PBMC samples used in this study. b. Dot plot of gene set enrichment analysis of most enriched immune pathways within Hallmark gene sets for each T/NK/ILC subpopulation in PBMCs. Mean expression of genes contained in each gene set within each cell type, separated into non-obese vs. obese groups, are indicated by colour gradients. P values are indicated by dot sizes. d. Dot plot of gene set enrichment analysis of most enriched immune pathways within Hallmark gene sets for each B/Plasma subpopulation in PBMCs. Mean expression of genes contained in each gene set within each cell type, separated into non-obese vs. obese groups, are indicated by colour gradients. P values are indicated by dot sizes.

**Figure S6.**
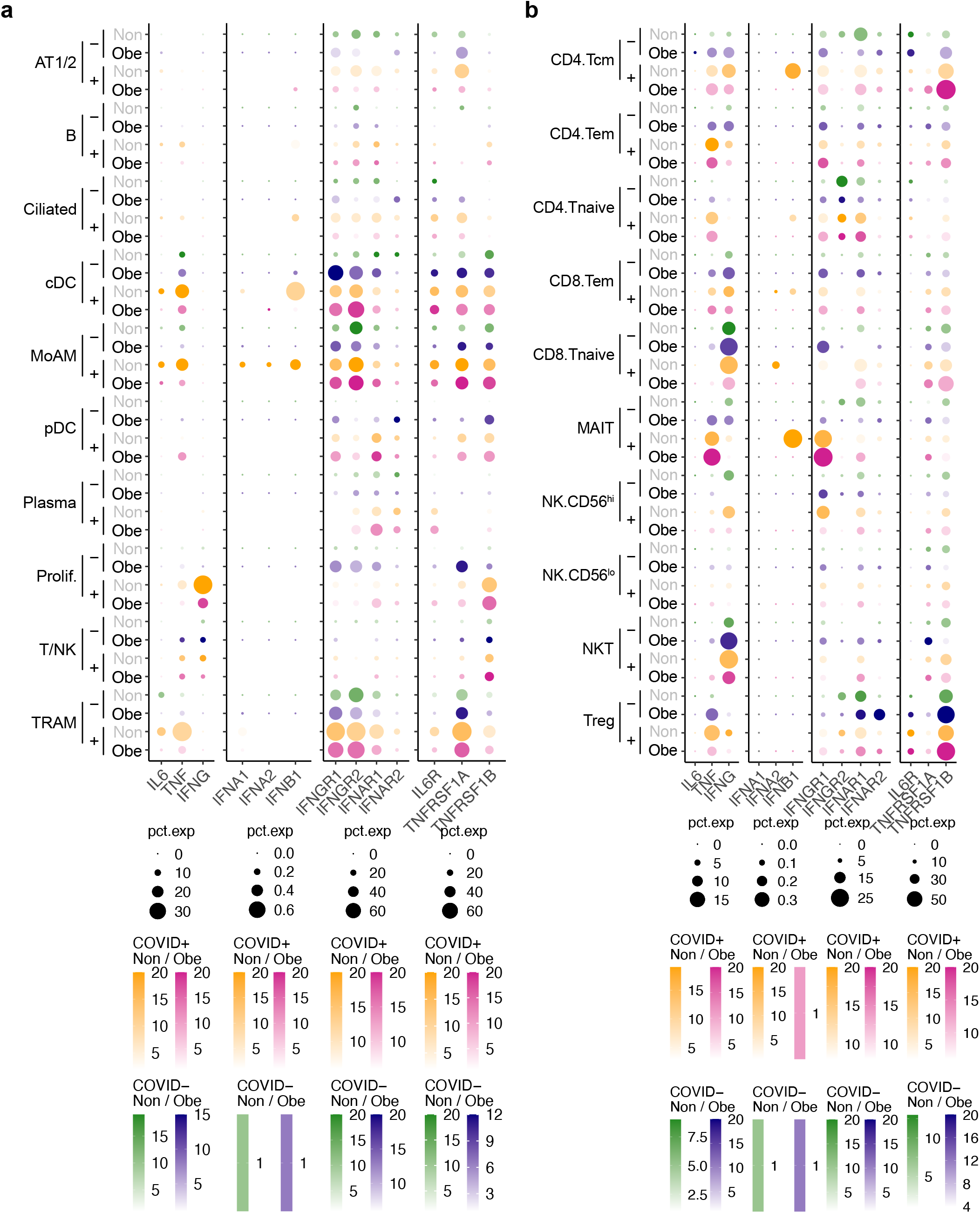
Expression patterns of the selected cytokines and their receptors in the BAL samples. a. Mean expression dot plots of transcripts for the selected cytokines and their receptors in each major population in the BAL samples. Expression levels in each case are indicated by distinct colour gradients (Green: Non-obese without COVID-19; Yellow: Non-obese with COVID-19; Purple: Obese without COVID-19; Magenta: Obese with COVID-19). Expression percentages are indicated by dot sizes. b. Mean expression dot plots of transcripts for the selected cytokines in each T/NK subpopulation in the BAL samples. Expression levels in each case are indicated by distinct colour gradients (Green: Non-obese without COVID-19; Yellow: Non-obese with COVID-19; Purple: Obese without COVID-19; Magenta: Obese with COVID-19). Expression percentages are indicated by dot sizes.

**Figure S7.**
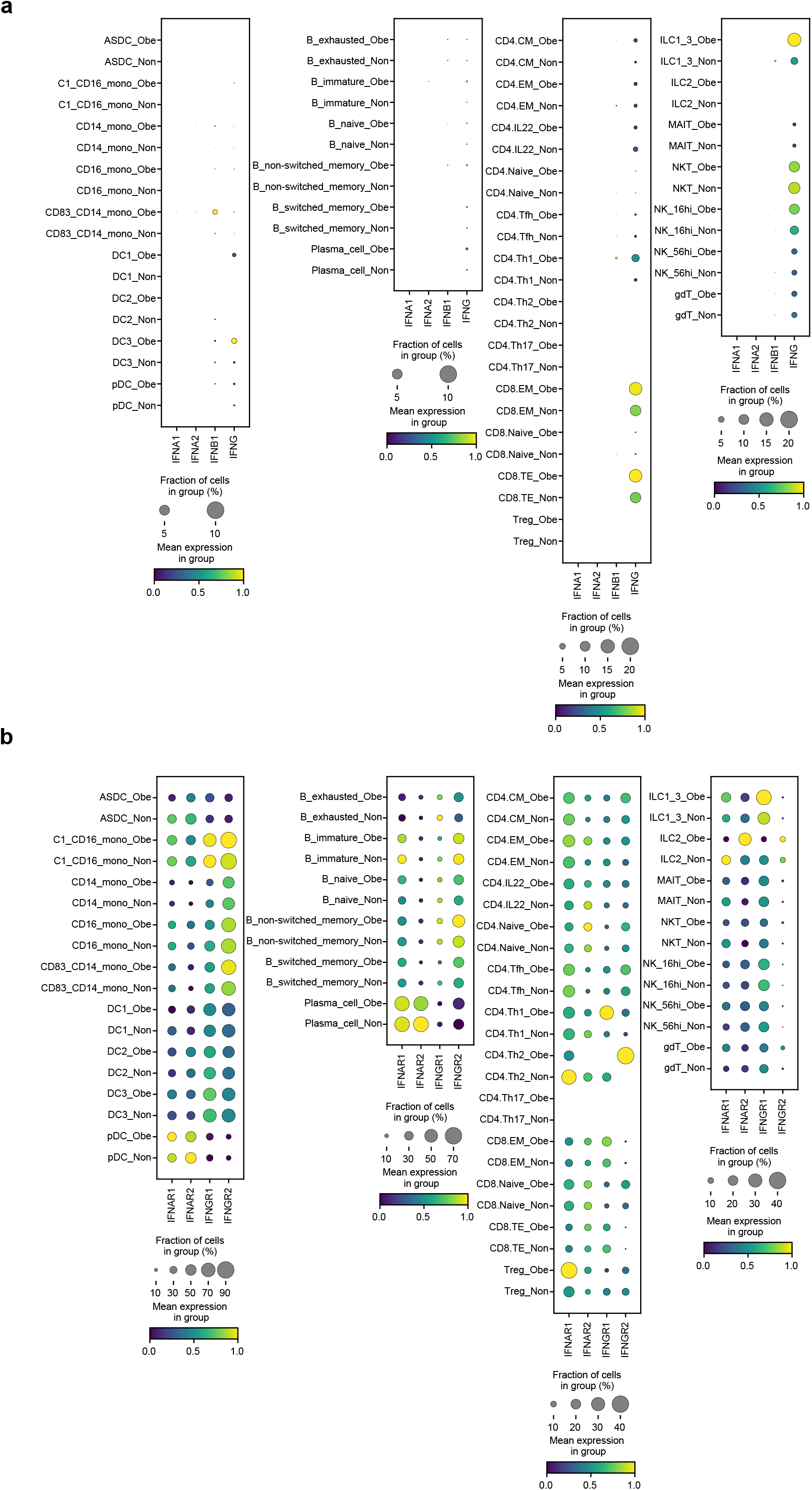
Expression patterns of the selected cytokines and their receptors in PBMCs. a. Mean expression dot plots for IFNA1, IFNA2, IFNB and IFNG in PBMC cell clusters from Stephenson et al. 2021^22^ grouped according to BMI status. Size of circles correspond to fraction of cells expressing each gene and increasing gradient from purple to yellow corresponds to increasing mean expression value (standardized to 0 to 1 per gene). b. Mean expression dot plots for IFNAR1, IFNAR2, IFNGR1 and IFNGR2 in PBMC cell clusters from^22^ grouped according to BMI status. Size of circles correspond to fraction of cells expressing each gene and increasing gradient from purple to yellow corresponds to increasing mean expression value (standardized to 0 to 1 per gene).

**Figure S8.**
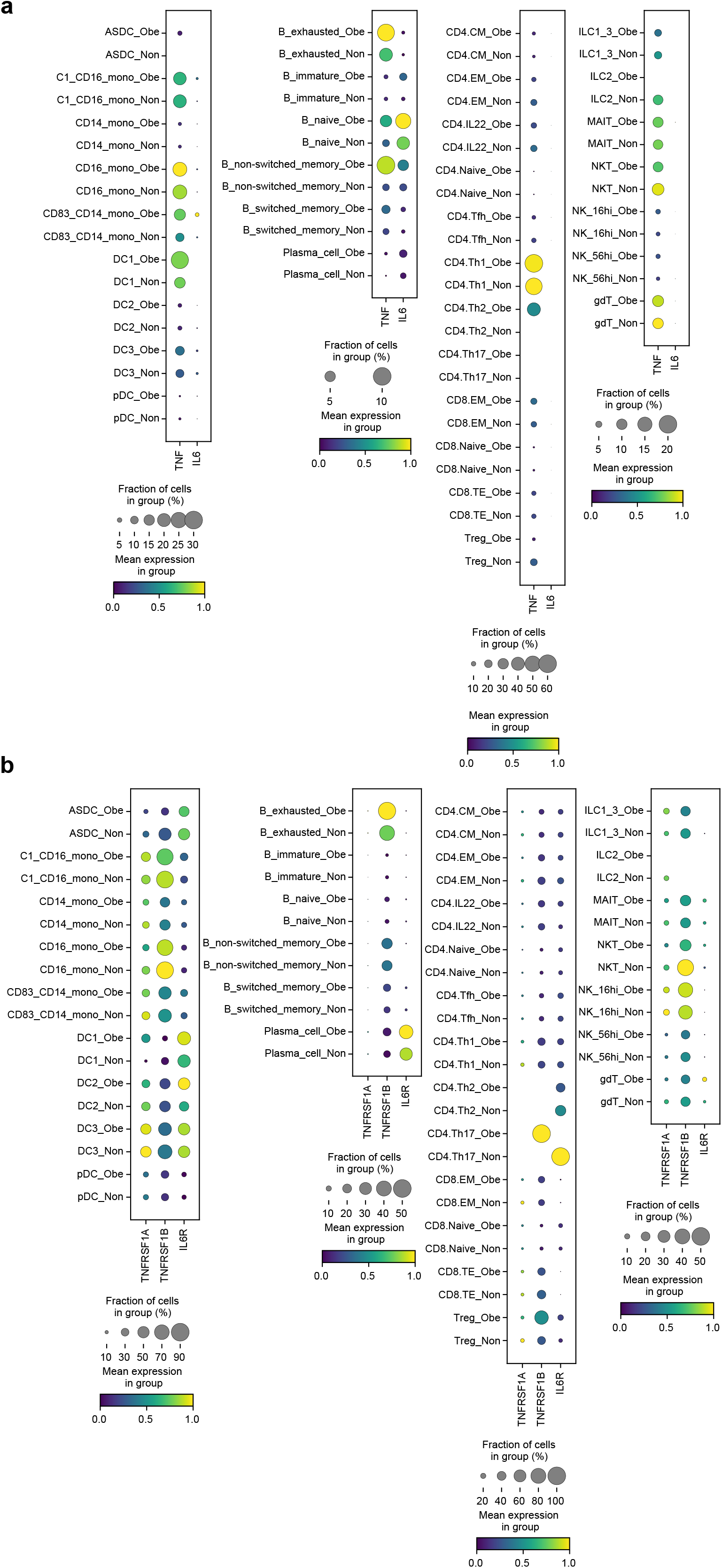
Expression patterns of the selected cytokines and their receptors in PBMCs. a. Mean expression dot plots for TNF and IL6 in PBMC cell clusters from Stephenson et al. 2021 ^22^ grouped according to BMI status. Size of circles correspond to fraction of cells expressing each gene and increasing gradient from purple to yellow corresponds to increasing mean expression value (standardized to 0 to 1 per gene). b. Mean expression dot plots for TNFRSF1A, TNFRSF1B and IL6R in PBMC cell clusters from^22^ grouped according to BMI status. Size of circles correspond to fraction of cells expressing each gene and increasing gradient from purple to yellow corresponds to increasing mean expression value (standardized to 0 to 1 per gene).

## Methods

### Patient recruitment and consent

The BAL samples from UCAM were collected in the Addenbrookes Hospital Intensive Care Unit under our discard lavage protocol. The use of discard samples surplus to that required for clinical testing, and anonymised data review were conducted under the consent waiver granted by Leeds West NHS Research Ethics Committee (ref: 20/YH/0152). Inclusion criteria were ‘adult (age >16) patients admitted to ICU for mechanical ventilation undergoing bronchoalveolar lavage for the investigation of suspected pneumonia’. Exclusion criteria were non-ventilated patients, age <16 years, and patients with restricted access to notes. For blood samples, ethical approval was obtained from the East of England – Cambridge Central Research Ethics Committee (“NIHR BioResource” REC ref 17/EE/0025, and “Genetic variation AND Altered Leukocyte Function in health and disease - GANDALF” REC ref 08/H0308/176). All participants provided informed consent. For paediatric nasal brushings, BMI data was obtained (where possible) on patients included in the UK Cohort of Yoshida *et al.* 2021 ^22^. Details of consent and methodology are found here.

### Isolation of the cells from BALF

Samples of 5-20ml BAL were collected and processed under BSL3 conditions. If necessary, the samples were filtered prior to processing by passing through a 100μm cell strainer to remove large mucus aggregates. Samples were subsequently topped up with PBS to 50ml and centrifuged at 400xg for 5 min. The supernatant was removed, and the cells were resuspended in 100μl of PBS. 20μl of Human fc block was added (Milteni) followed by 10μl of a custom TotalSeq -C Human cocktail. Cells were stained for 30min, topped up with PBS and washed as before.

### Single-cell encapsulation, library preparation, and sequencing

Cells were resuspended in 50μl, counted, and loaded on Chromium Chip A (10x Genomics, 5’v3) for cell encapsulation. cDNA libraries were prepared per manufacturer’s recommendation. After quality checks with Bioanalyzer (Agilent; 2100), the libraries were pooled and sequenced with NovaSeq 6000.

### Isolation of PBMCs and granulocytes

PBMC and granulocytes were isolated by discontinuous density gradient centrifugation in 60% and 80% Percoll at 800g for 15 minutes. PBMC and granulocyte layers were taken off and washed separately in PBS at 300xg for 10 minutes. Cells were counted and 1-3×10^6^ cells from each of the PBMC and granulocyte layers were separately stained for flow cytometry.

### Flow cytometry

After aliquots were taken for single-cell RNA sequencing, remaining BAL fluid cells, PBMC and granulocytes were blocked with human FcR block (Miltenyi Biotech, Bisley, UK) and incubated with antibodies (see table) for 30 minutes at 4°C, then washed in PBS and resuspended in FACS fix. Samples were processed on a Fortessa flow cytometer (Becton Dickinson, Basel, Switzerland) and data analysed using Flowjo version 10.

**Table 1.**
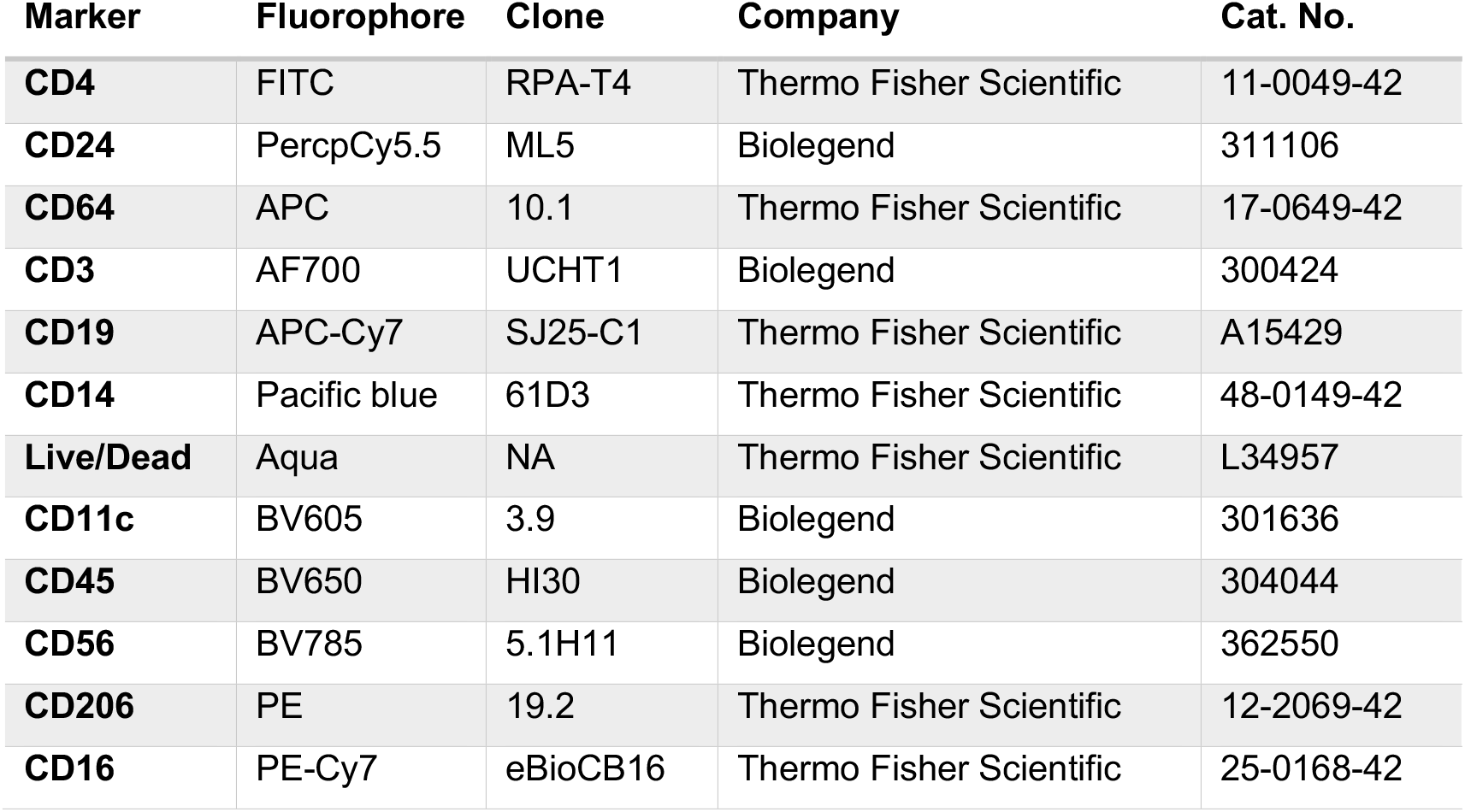
List of antibodies used in flow cytometry

### Public datasets

We have included two public single-cell RNA-seq datasets from Liao et al. (2020)^18^ and Grant et al. (2021)^19^ for multi-sample integration. The data were acquired from Gene Expression Omnibus (GEO) database under accession codes GSE145926 and GSE155249, respectively. The additional healthy control BAL sample (Morse et al., 2019; GSE128033)^30^ analyzed in Liao et al. (2020)^18^ has also been included.

### Single-cell RNA-seq data alignment and multi-sample integration

Data were processed using the Cell Ranger 3.1.0 pipeline (10x Genomics). The generated count tables with UMI counts > 1000, gene number > 200, and mitochondrial genes < 10% were analysed using the Seurat^31^ software package (4.0.4) in R (4.0), ambient RNA was corrected with SoupX^32^ (1.5.2), doublets were detected with DoubletFinder^33^ (2.0.3) and removed, and multi-sample integration was performed with harmony^34^ (1.0) to remove batch effects across different patients. In parameter settings, the first 50 dimensions of principal-component analysis (PCA) were used, and the cells were clustered using the FindClusters function with a resolution of 0.5. Uniform Manifold Approximation and Projection (UMAP)^35^ v(0.5.2) was used for visualizing the data.

### Differential gene expression analysis

Wilcox test in FindAllMarkers or FindMarkers function in Seurat was used to compare the differential gene expression across major or minor cell types or within each subpopulation of structural cells, macrophages, T/NK cells, B/Plasma cells, or dendritic cells between non-obese and obese cases, respectively.

### Re-integration of structural cells, macrophages, T / NK cells, and B cells

Major cell types including structural cells, macrophages, T / NK cells, and B cells were re-clustered with Seurat (4.0.4), followed by removal of batch effect across different patients with harmony (1.0). Cells were clustered using the FindClusters function with resolutions between 0.4-1.0 (structural cells: 0.5; myeloid cells: 0.4; T cells: 1.0; B cells: 1.0).

### Gene set enrichment analysis (GSEA)

clusterProfiler^36^ (3.18.1) was used to perform GSEA. Briefly, genes from each of the subpopulations were ranked in a descending order of their expression levels between non-obese and obese cases by using FindMarker function. Those with log fold changes > 0.5 or < −0.5 were selected for GSEA using compareCluster function. Hallmark gene sets in msigdbr (7.4.1) and enricher function in clusterProfiler were used for gene functional annotation.

### Calculation of the cytokine module score

The module score was calculated using AddModuleScore function in Seurat with the cytokine and chemokine gene set from KEGG pathway.

### Ligand-receptor analysis

Ligand-receptor analysis was performed with CellPhoneDB^37^ and was visualized with ktplots using plot_cpdb function. Briefly, the normalized counts and meta data extracted from Seurat objects were applied for the statistical analysis from CellPhoneDB in python 3.8.8. The resulting p values and means were then filtered and visualized with ktplots.

### Visualization

Plotting was performed using ggplot2 (3.3.5). Heatmap was generated using pheatmap (1.0.12). Figure layouts were edited in Affinity Designer (1.10.0).

### Statistics

Statistical analysis was performed using base R (4.0) with tidyverse (1.3.0). Wilcoxon tests were performed in FigS1a using stat_compare_means function in ggpubr with ‘wilcox.test’ indicated in method parameter

